# A human mitochondrial isoform of TRPV1 regulates intracellular Ca^2+^ simultaneously with mitochondrial thermolysis

**DOI:** 10.1101/2024.09.25.614940

**Authors:** Florian Beignon, Sylvie Ducreux, Léa Tuifua, Yannick Le Dantec, Morgane LeMao, David Goudenège, Arnaud Chevrollier, Salim Khiati, Hélène Tricoire-Leignel, Naig Gueguen, César Mattei, Guy Lenaers

**Affiliations:** Univ Angers, MitoLab, Unité MITOVASC, UMR CNRS 6015, INSERM U1083, SFR ICAT, Angers, France; Univ Claude Bernard Lyon1, CarMeN Laboratory, INSERM, INRA, 69500 Bron, France; Univ Angers, CarMe, Unité MITOVASC, UMR CNRS 6015, INSERM U1083, SFR ICAT, Angers, France; Service de Biochimie et Biologie Moléculaire, CHU d’Angers, Angers, France; Service de Neurologie, CHU d’Angers, Angers, France

**Author notes:** Ifremer, IRSI, SeBiMER Service de Bioinformatique de l’Ifremer, F-29280 Plouzané, France.

## Abstract

Mitochondria are the cornerstones of cellular and body thermogenesis, with an inner temperature possibly reaching 50°C. Here, we report the identification of a human Transient Receptor Potential Vanilloid 1 alternative isoform located in mitochondria. This isoform, which we have termed mitoTRPV1, acts as a thermostat to restrict the mitochondrial temperature. The mitoTRPV1 open reading frame overlaps *TRPV1* exons 1 and 2 and intron 2 in a +1 frame, encoding for a predicted 150 amino-acid N-terminal mitochondrial targeting sequence (MTS) conserved amongst mammalian species, followed by the 687 amino acids of TRPV1 C-terminal. This ORF is ubiquitously expressed in most human organs, underscoring its broad relevance. The deduced MTS, conserved among mammalian species, effectively addresses this TRPV1 isoform to the mitochondrial inner membrane. Our experiments, using heterologous wild-type and mutated mitoTRPV1 expression, combined with Ca^2+^ imaging, mitochondrial temperature and oxygraphy measurements, disclosed that mitoTRPV1 activation induces Ca^2+^ efflux and mitochondrial cooling, without modification of mitochondrial respiration and ATP production. Notably, the loss of function mitoTRPV1-G684V isoform, responsible for exertional heat stroke predisposition in humans, abolished mitochondrial Ca^2+^ efflux and cooling. These findings reveal a new thermolysis function for TRPV1 in preventing mitochondrial overwarming while not affecting the OXPHOS efficiency. They also highlight the potential implications of mitoTRPV1 in human diseases related to temperature dysregulation.

## Introduction

Mammals are homeothermic species that balance thermogenesis and thermolysis to adapt body temperature in changing environmental conditions, a critical process, as variations of plus or minus 5°C lead to morbidity and lethality^1,2^. Thermogenesis relies on heat generated by mitochondrial oxidation process, promoting mitochondria as cell radiators^3^. In line with this concept, the mitochondrial temperature in cells can be up to 15°C above that of their external environment, thus reaching 50°C, a temperature at which the enzymatic activity of most of the oxidative phosphorylation (OXPHOS) respiratory chain complexes are most efficient^4,5^. This hypothesis is, however, a matter of debate^6^ as the mechanisms sensing and monitoring mitochondrial maximal temperature are totally unknown. Thus, today, we have a limited understanding of mitochondrial thermogenesis control and its contribution to homeothermy in mammals, although the Ca^2+^ channel TRPV4 was demonstrated to localize to a subpart of the mitochondrial network where it promotes Ca^2+^ entry and activates thermogenesis at temperature below 37°C^7^. To gain insights into this topic, we hypothesized that mitochondria must include a thermo-sensor to modulate thermogenesis and prevent deleterious effects associated with hyperthermia.

Thermo-TRPs are Transient Receptor Potential (TRP) ion channels activated by heat (TRPV1-4, TRPM2-5) or by cold (TRPA1, TRPM8 and TRPC5) and act as thermosensors in the peripheral nervous system to detect environmental temperatures^8^. Most TRPs are localized at the plasma membrane, or in intracellular organelles, including mitochondria, but none of them has ever been proposed to act as a thermostat limiting mitochondrial warming.

TRPV1 is a polymodal cation channel activated by temperatures above 43°C, by capsaicin (CAP) - the pungent “burning” component of red chili peppers - and by resiniferatoxin (RTX), its most specific agonist^9,10^. TRPV1 is implicated in the noxious sensing of thermal and chemical stimuli, which activate somatosensory neurons responsible for acute and inflammatory pains^11^. Consequently, TRPV1 activation in primary afferent sensory neurons lowers body temperature by promoting vasodilation and sweating^12,13^, while conversely, its inhibition displays antalgic actions and induces deleterious hyperthermia symptoms^14^.

In line with these data, *TRPV1* variants were associated with a predisposition to acute life-threatening hyperthermia, both in individuals facing exertional heat strokes and in patients treated with halogenated anaesthetics^15,16^. In all cases, syndromes were associated with a failure to counteract hyperthermia with dramatic consequences on brain, muscle, liver, and kidney integrity^17,18^. Additional *TRPV1* variants abrogating TRPV1 activity or partial deletions of *TRPV1* promoter were identified in patients presenting heat hyposensitivity, delayed response to noxious heat, inflammatory hyperalgesia, and insensitivity to CAP^19,20^. These clinical data demonstrate that TRPV1 is fundamental to respond to thermal and noxious stresses and adapt body temperature, while *Trpv1-*KO mouse models only illustrated altered responses to noxious stimuli without any thermoregulation defect^21^. Beyond these roles, alternative TRPV1 expression and localization led to novel TRPV1 functions in neuronal and non-neuronal cells^22^. Indeed, aside to its embedment in the plasma membrane, TRPV1 was also localized in intracellular compartments, as the sarcoplasmic reticulum membrane of skeletal muscle, acting as a Ca^2+^ leak channel^23^. It is also expressed in the mitochondria of cardiomyocytes, where it stimulates ATP synthesis^24^, in microglial cells to promote cell migration^25^, and in endothelial cells contributing to diabetes onset^26^. Nevertheless, the role of TRPV1 in intracellular temperature sensing has yet never been reported. Here, we describe and characterize a novel mammal-specific TRPV1 mRNA variant encoding an isoform located in the mitochondrial inner membrane (hereafter called mitoTRPV1) and its involvement in mitochondrial Ca^2+^ homeostasis, thermogenesis, and respiration, comparatively to the conventional TRPV1 isoform located at the plasma membrane (hereafter called pmTRPV1).

## Results

### mitoTRPV1 is encoded by a novel atypical ORF

To identify a mitochondrial “hot” thermostat, we screened databases for TRP variants, including a potential mitochondrial targeting sequence (MTS) using the MitoProt-II prediction software V1.101. This process identified a *TRPV1* cDNA (accession number DQ177332.1 (NCBI) and ENST00000574085.5 (Ensembl)) starting by an alternate ATG initiation codon in exon1, generating an overlapping +1 ORF encompassing exon1 3’-end, the complete exon2, the 70 nt-long intron2, followed by *TRPV1* consensual coding frame from exon3 to exon16 (Fig. 1a). Translation of the nucleotide sequence leads to a predicted 150 amino acid (aa) MTS sequence at the N-terminal domain (0.9968 probability according to MitoProt-II and a predicted cleavage site at position 138) followed by the 687 aa of TRPV1 C-terminal (Fig. 1b). This isoform was called mitoTRPV1.

**Figure 1.**
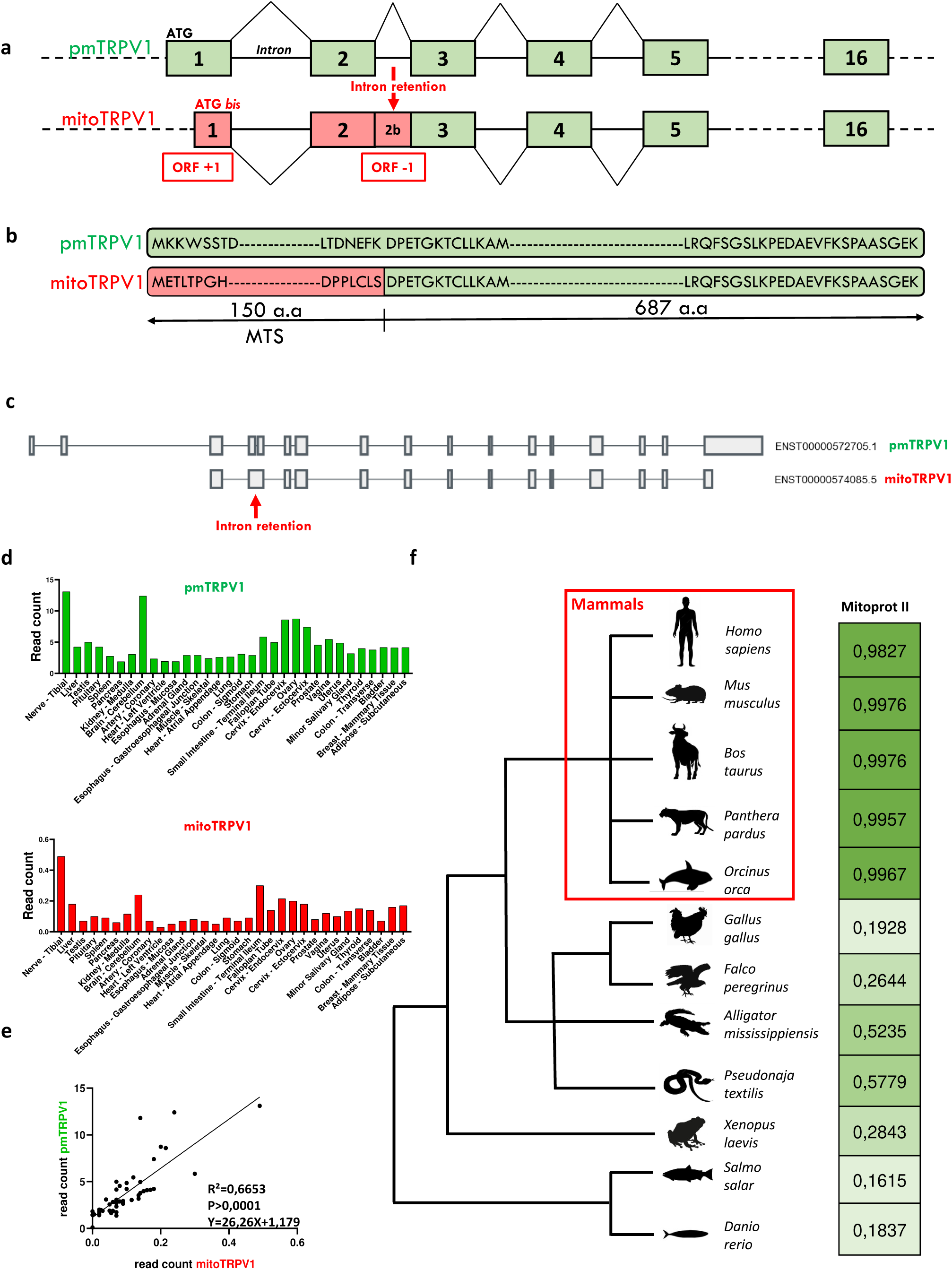
Identification of mitoTRPV1 cDNA and protein sequence. **a,** Open reading frame organization of pmTRPV1 and mitoTRPV1 variants. MitoTRPV1 ATG initiation codon lies in the first coding exon in a +1 frame and generates an overlapping ORF encompassing also exon2 and the 70 nt long intron2, which retention restores the consensual TRPV1 ORF. **b,** Comparison of the protein structure of the two TRPV1 isoforms, the one located in the plasma membrane (pmTRPV1) on top and the one with a mitochondrial targeting sequence (MTS in pink, amino acids 1–150) (mitoTRPV1) on the bottom. **c,** Structures of TRPV1 coding sequence of pmTRPV1 and mitoTRPV1 with intron 2 retention reported in the GTEx Portal database. **d,** Quantitative tissue expression of the pmTRPV1 (green) and mitoTRPV1 (red) mRNA variants in the GTEx Portal database. **e,** Ratio (bottom) between mitoTPRV1 and pmTRPV1 expression, inferred from the histograms and values from d. **f,** Phylogenetic prediction for the presence of an MTS in an alternate ORF in TRPV1 gene from the major vertebrate species lineages, using the Mitoprot II software. Mammals are boxed in red.

Browsing the GTEx portal database (https://www.gtexportal.org/home/, accessed on 18 September 2024) confirmed the existence of *mitoTRPV1* mRNA with the intron2 retention between exon2 and 3 (Fig. 1c). It disclosed its ubiquitous expression in 32 human tissues, paralleling that of pmTRPV1 mRNA (R²=0.6653), although with an abundance 26 times lower than that of the conventional pmTRPV1 (Fig. 1d-e).

To extend this discovery, we screened for the presence of an MTS in *TRPV1* genomic sequence from other vertebrate animal species using the MitoProt-II program, considering all possible initiation codon extensions. We evidenced that the probability of generating an MTS is close to 1 in mammal species, while this score was significantly lower in homeothermic birds or poikilotherm reptiles and fishes (Fig. 1f).

### MitoTRPV1 is located in the mitochondrial inner membrane

To challenge the predictive *in silico* value of mitoTRPV1 MTS and its functional relevance, we cloned GFP fusion proteins, including the whole or parts of the mitoTRPV1 sequence and expressed them in MCF7 cells. Compared to the Mitotracker fluorescence, expression of the GFP alone or mitoTRPV1 deleted of the MTS sequence displayed a cytoplasmic localization. Conversely, the expression of mitoTRPV1 or its MTS alone in fusion with the GFP displayed a mitochondrial fluorescence superposed to that of the Mitotracker (Fig. 2a). This result indicates that mitoTRPV1 MTS is required and sufficient to drive mitoTRPV1 to mitochondria. Further Western blot analysis of a mitochondrial fraction of HEK293 cells expressing mitoTRPV1-GFP confirmed the exclusive mitochondrial localization of mitoTRPV1 (Fig. 2b). To address the mitochondrial sub-localization of mitoTRPV1, we first excluded that mitoTRPV1 is located in MAMs (Extended Data Fig. 2), then we performed a Proteinase-K protection assay on mitochondria purified from HEK293 cells expressing mitoTRPV1-6His. Without treatment, mitoTRPV1 appears as two bands, one corresponding to the full-length protein, the other as a shorter isoform probably cleaved of its MTS (Fig. 2c, lane 1). Initial treatment with Proteinase-K in the absence of digitonin led to a second mitoTRPV1 cleavage, paralleling the degradation of TOM20 located at the outer membrane (Fig. 2c, lane 2), suggesting that either the N-ter (after the MTS cleavage) or the C-ter end of mitoTRPV1 lays in the intermembrane space (IMS). The addition of increasing digitonin concentrations led to a mitoTRPV1 protection pattern similar to that of TIM23, suggesting that the core of the mitoTRPV1 protein is embedded in the mitochondrial inner membrane (Fig. 2c, lanes 3 to 6). This result is also supported by the “DeepMito” tool (http://busca.biocomp.unibo.it/deepmito/, accessed on 18 September 2024) that predicts mitoTRPV1 localization in the mitochondrial inner membrane with a score of 0.73. The presence of an MTS indicates that mitoTRPV1 is unequivocally a mitochondrial protein, with its core-channel domain firmly embedded in the inner membrane.

**Figure 2.**
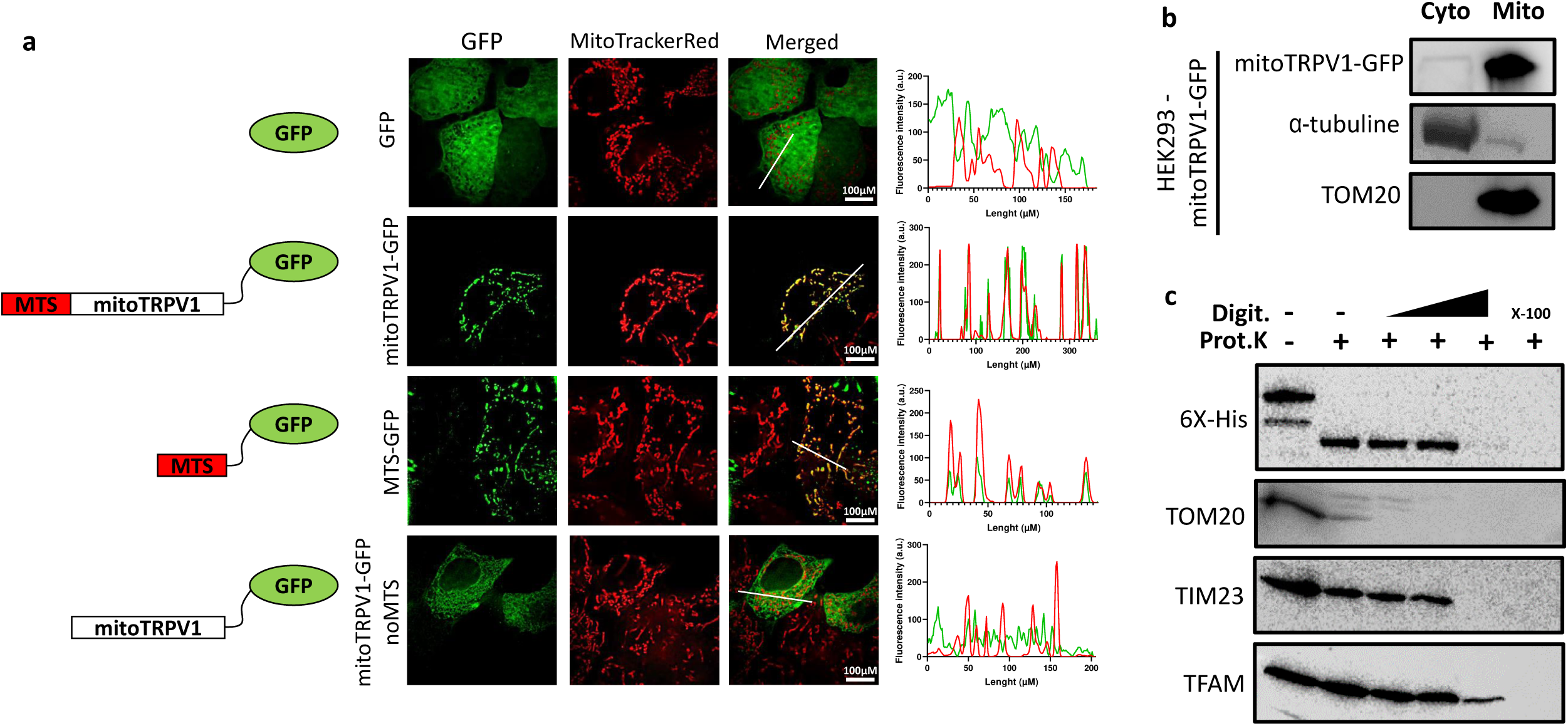
Mitochondrial localization of mitoTRPV1. **a,** Co-localization analysis of GFP (green) and Mitotracker (red) fluorescence in MCF-7 cells expressing GFP alone (1^st^ line), mitoTRPV1-GFP (2^nd^ line), MTS-GFP (3^rd^ line) and mitoTRPV1-GFP deleted from the MTS (4^th^ line), together with the histogram transection levels of green and red fluorescences. **b**, Western blot analysis of cytoplasmic (left) and mitochondrial (right) fractions of HEK293 cells expressing mitoTRPV1. **c,** Proteinase K protection assay of a mitochondrial fraction from cells expressing a 6-His-tagged mitoTRPV1 isoform. Column 1 corresponds to the untreated mitochondrial fraction. Columns 2 to 6 show the fraction treated with proteinase K without digitonin (column 2), with increasing concentrations of digitonin (column 3-5), or with Triton X100 (column 6). Representative Western blots obtained with antibodies against 6-His-tagged TOM20, TIM23, and TFAM.

### mitoTRPV1 promotes Ca^2+^ efflux from mitochondria

To assess mitoTRPV1 channel activity, Ca^2+^ variations were evaluated in HEK293 cells expressing either the mitoTRPV1 or the pmTRPV1 isoform, stimulated by RTX, using a FlexStation device to assess cellular populations or a dedicated fluorescent microscope to assess single transfected cells.

First, cytosolic Ca^2+^ variations ([Ca^2+^]_c_) were measured with Fura-2. The resting [Ca^2+^]_c_ was not modified by mitoTRPV1 expression but was increased by pmTRPV1 expression (Extended Data Fig. 3a). As expected, pmTRPV1 activation by RTX led to a solid dose-dependent increase in [Ca^2+^]_c_ due to a massive Ca^2+^ entry from the extracellular medium (Fig. 3a). Similarly, mitoTRPV1 activation led to a [Ca^2+^]_c_ increase, although lower compared to pmTRPV1 activation, and was fully inhibited by the TRPV1 antagonist capsazepine (Fig. 3b). EC_50_ values were 124.3 nM and 88.3 nM respectively for mitoTRPV1 and pmTRPV1 (Fig. 3c) and were independent of their respective expression level (Extended Data Fig. 1b). These results were confirmed by a single-cell imaging method (Fig. 3d-e), suggesting that upon activation by RTX, mitoTRPV1 promotes Ca^2+^ efflux from mitochondria, resulting in a [Ca^2+^]_c_ increase, with pharmacological properties similar to pmTRPV1 with respect to RTX activation and capsazepine inhibition.

**Figure 3.**
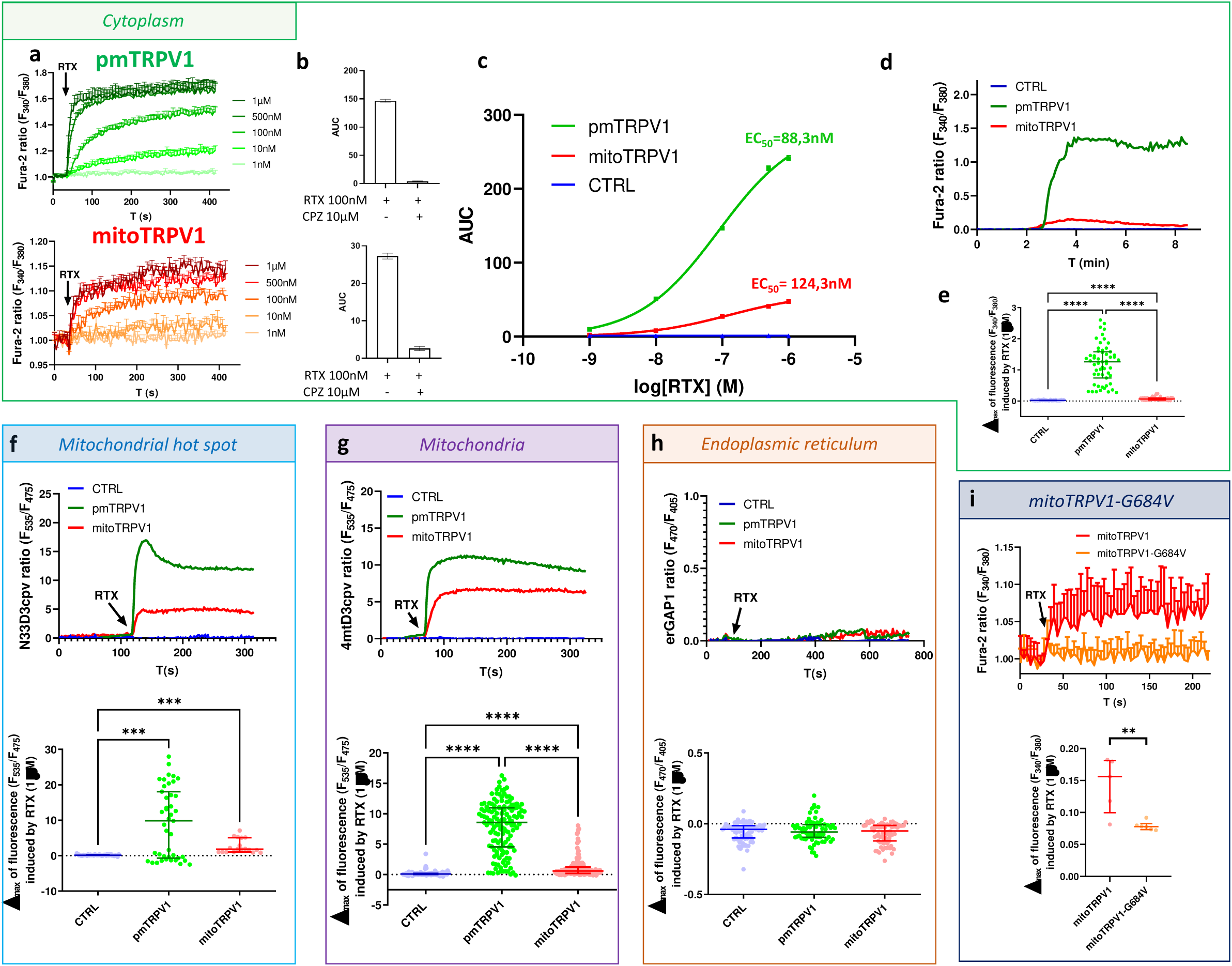
Activation of mitoTRPV1 drives Ca^2+^ flux from mitochondria. **a,** Time traces of normalized Fura-2 fluorescence ratio, reflecting the cytoplasmic Ca^2+^ concentration ([Ca^2+^]_c_), in HEK293 cells expressing pmTRPV1 (light to dark green) or mitoTRPV1 (orange to red) stimulated with increasing concentrations of RTX (from 1 nM to 1 µM). **b,** Inhibitory effect of capsazepine (CPZ, 10 µM) on RTX-induced [Ca^2+^]_c_ increase in HEK293 cells expressing pmTRPV1 (top) or mitoTRPV1 (down) (n=3). **c,** RTX dose-response curve of HEK293 cells expressing pmTRPV1 (green) or mitoTRPV1 (red) isoform and control cells (blue), inferred from the results described in **a,** and determination of the corresponding EC_50_ value using GraphPad Prism 8. **d**, Representative time traces of normalized Fura-2 fluorescence ratio, reflecting the [Ca^2+^]_c_ in HEK293 cells expressing pmTRPV1 (green) or mitoTRPV1 (red) isoform and control cells (blue) stimulated with RTX (1 µM), using single-cell fluorescence microscopy. **e,** Scatter plots representing the corresponding cell-by-cell maximal amplitude of RTX-response (data collected in **d**: CTRL, n=101; pmTRPV1, n=54; mitoTRPV1, n=53). **f,** Top: Representative time traces of normalized N33D3cpv fluorescence ratio, reflecting the mitochondrial hot spot Ca^2+^ concentration ([Ca^2+^]_hot spot_), in HEK293 cells expressing pmTRPV1 (green) or mitoTRPV1 (red) isoform and control cells (blue) stimulated with RTX (1 µM). Bottom: scatter plots representing the corresponding cell-by-cell maximal amplitude of RTX-response (CTRL, n=38; pmTRPV1, n=43; mitoTRPV1, n=18). **g,** Top: Representative time traces of normalized 4mtD3cpv fluorescence ratio, reflecting the mitochondrial Ca^2+^ concentration ([Ca^2+^]_mito_), in HEK293 cells expressing pmTRPV1 (green) or mitoTRPV1 (red) isoform and control cells (blue) stimulated with RTX (1 µM). Bottom: scatter plots representing the corresponding cell-by-cell maximal amplitude of RTX-response (CTRL, n=92; pmTRPV1, n=141; mitoTRPV1, n=116). **h,** Top: Representative time traces of normalized erGAP1 fluorescence ratio, reflecting the endoplasmic reticulum Ca^2+^ concentration([Ca^2+^]_ER_), in HEK293 cells expressing pmTRPV1 (green) or mitoTRPV1 (red) isoform, and control cells (blue) stimulated with RTX (1 µM). Bottom: scatter plots representing the corresponding cell-by-cell maximal amplitude of RTX-response (CTRL, n=102; pmTRPV1, n=75; mitoTRPV1, n=72). **i,** Top: Time traces of normalized Fura-2 fluorescence ratio, reflecting [Ca^2+^]_c_, in HEK293 cells expressing mitoTRPV1 (red) or mitoTRPV1-G684V (orange) isoform stimulated with RTX (1 µM), using single-cell fluorescence microscopy. Bottom: scatter plots representing the corresponding cell-by-cell maximal amplitude of RTX-response (mitoTRPV1, n=5; mitoTRPV1-G684V, n=6). Data are from at least three independent experiments. P-value correspondences *: p<0.05, **: p<0.01, ***: p<0.001, **** p<0.0001

To gain insights into the intracellular Ca^2+^ modifications induced by mitoTRPV1 activation, mitochondrial surface hot spot Ca^2+^ variations ([Ca^2+^]_hot spot_) were measured with N33D3cpv, a Ca^2+^ sensor embedded in the mitochondrial outer membrane. mitoTRPV1 overexpression did not modify the resting [Ca^2+^]_hot spot_, while pmTRPV1 overexpression increased the resting [Ca^2+^]_hot spot_ (Extended Data 3b). The activation of pmTRPV1 with RTX further increased [Ca^2+^]_hot spot_ due to massive Ca^2+^ influx from the extracellular medium into the cytosol. Similarly, mitoTRPV1 activation increased [Ca^2+^]_hot spot_, although to a lower extent, suggesting that mitoTRPV1 promotes Ca^2+^ exit from mitochondria (Fig. 3f), thus confirming data from Fig. 3d-e.

Complementarily, we assessed mitochondrial matrix Ca^2+^ variations ([Ca^2+^]_mito_) using the 4mtD3cpv, a mitochondrial matrix Ca^2+^ sensor. MitoTRPV1 overexpression did not modify the resting [Ca^2+^]_mito_, while pmTRPV1 overexpression increased the resting [Ca^2+^]_mito_ (Extended Data Fig. 3c). As expected, pmTRPV1 activation with RTX strongly increased [Ca^2+^]_mito_ associated with cytoplasmic Ca^2+^ buffering by the mitochondrial matrix. Unexpectedly, mitoTRPV1 activation with RTX also led to a significant faint mitochondrial Ca^2+^ increase in cells (Fig. 3g). Taken together, these results show that mitoTRPV1 triggers a Ca^2+^ efflux from mitochondria paralleled by a Ca^2+^ reentry. Of note, measurements of endoplasmic reticulum (ER) Ca^2+^ variations ([Ca^2+^]_ER_) using the erGAP1 sensor did not evidence changes in [Ca^2+^]_ER_ resting condition (Extended Data Fig. 3d) or after activation of mitoTRPV1 or pmTRPV1 with RTX (Fig. 3f), indicating that mitoTRPV1 does not significantly affect ER Ca^2+^ load.

Ca^2+^ measurements were repeated with the mitoTRPV1 variant including the G684V amino-acid change, a loss of function mutation identified in exertional heat stroke patients (EHS)^16^. As expected, activation of mitoTRPV1-G684V with RTX (Fig. 3i) or CAP (Extended Data Fig. 5) did not modify the [Ca^2+^]_c_, while we confirmed that its expression level (Fig. S1c) and its localization into mitochondria (Extended Data Fig. 4) were identical to that of wild-type mitoTRPV1. Thus, mitoTRPV1-G684V does not induce Ca^2+^ efflux from mitochondria.

### mitoTRPV1 reduces mitochondrial temperature without altering cellular respiration

To assess the role of mitoTRPV1 in controlling mitochondrial temperature and respiration, we monitored MTY fluorescence simultaneously to oxygen consumption (Fig. 4a) in HEK293 cells expressing mitoTRPV1 or mitoTRPV1-G684V, activated or not with RTX. As previously reported for control HEK cells^4^, the mitochondrial temperature reached 51°C (Fig. 4b and 4d), while the addition of RTX led to a non-significant decrease to 49°C (Fig. 4b and 4e). In the same experimental conditions, we observed that mitoTRPV1 expression reduced the mitochondrial temperature to 44°C (Fig. 4b and 4f) and further to 40°C in the presence of RTX (Fig. 4b and 4g). In comparison, the expression of mitoTRPV1-G684V (Fig. 4b and 4h) resulted in a mitochondrial temperature similar to that of wild-type cells, suggesting that mitochondrial temperature downregulation depends on the Ca^2+^ efflux. Importantly, in all experimental conditions tested, the cell oxygen consumption rates were identical (Fig. 4c and 4d-h), witnessing that the mitochondrial temperature restriction is not associated to a mitochondrial respiration decrease.

**Figure 4.**
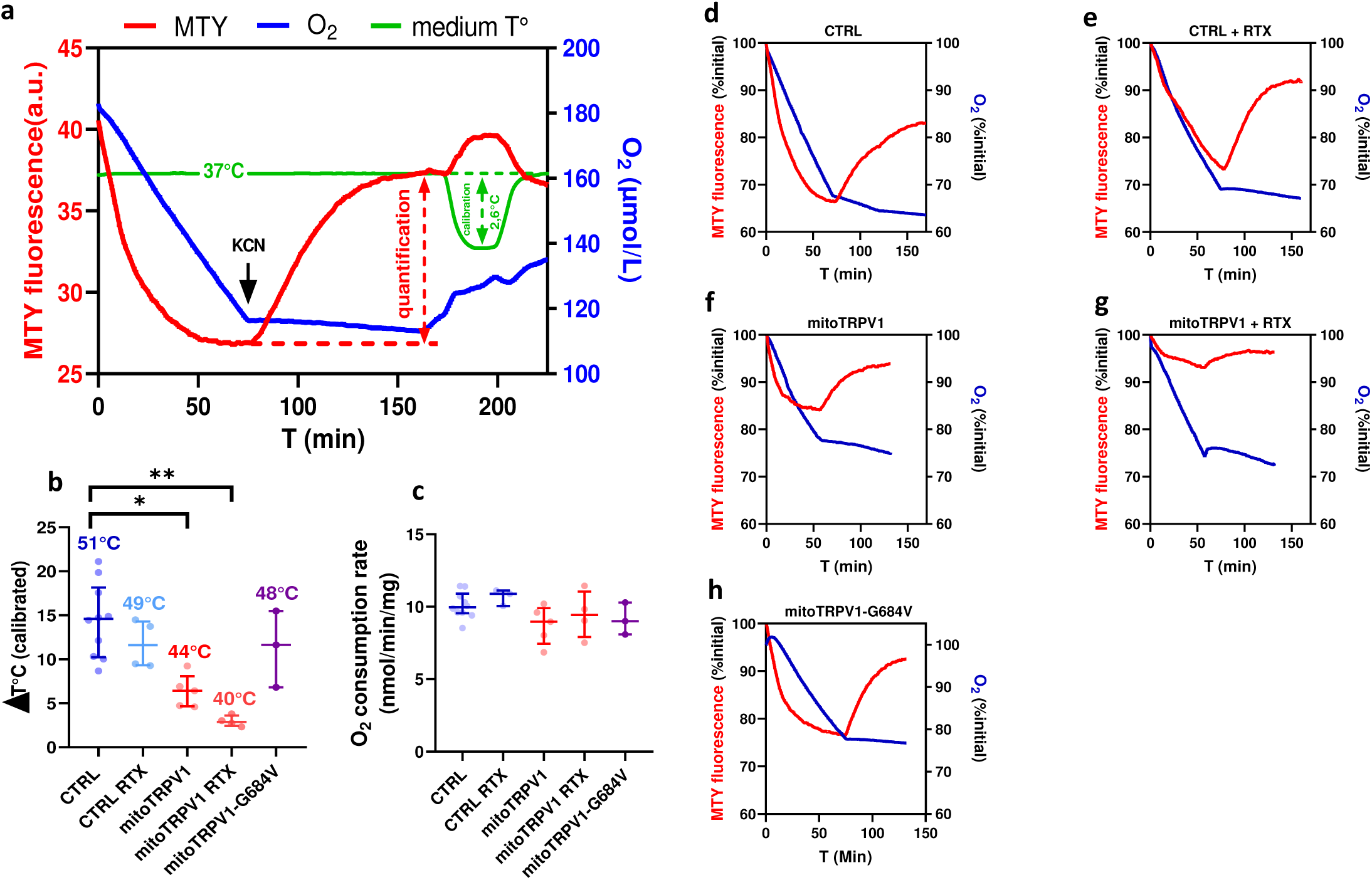
mitoTRPV1 expression and activation reduce mitochondrial temperature without affecting cell respiration. **a,** Experimental design: a representative time trace of MTY fluorescence (blue), O2 consumption (red), and cell medium temperature (green) during the experience. Black arrow: KCN addition. **b,** Mitochondrial temperature (ΔT°C) inferred from MTY fluorescence measurements and calibration from experiments **d-h**. **c,** Quantification of 0_2_ consumption (nmol/min/mg) from experiments **d-h**. **d and e,** Representative time traces of MTY fluorescence and O^2^ consumption of CTRL HEK293 cells without (d) or with RTX (1 µM) (e). **f and g,** Representative time traces of MTY fluorescence and O^2^ consumption of HEK293 cells expressing mitoTRPV1 without (f) or with RTX (1 µM) (g). **h,** Representative time traces of MTY fluorescence and O^2^ consumption of HEK293 cells expressing mitoTRPV1-G684V. Data are from at least three independent experiments. P-value correspondences *: p<0.05, **: p<0.01, ***: p<0.001, **** p<0.0001

To confirm the absence of effect of mitoTRPV1 on oxygen consumption, we used the Seahorse XF96 device (Fig. 5a) to monitor mitochondrial respiration in cells expressing mitoTRPV1 or pmTRPV1 in the absence or presence of RTX. Expression of pmTRPV1 increased basal respiration (R), respiration linked to ATP synthesis (R-O), maximal respiration (Fmax), respiration linked to proton leak (O), the proportion of respiration linked to proton leak compared to maximal respiratory capacity (O/F) and reduced spare respiratory capacity (R/F ratio) (Fig. 5b-5h), comparatively to untransfected cells. In addition, the ATP levels in mitochondria measured by flow cytometry (Extended Data Fig. 7) using ATeam sensor (Fig. 5i) were also reduced in pmTRPV1-expressing cells. Conversely, expression of mitoTRPV1 did not significantly change respiration parameters (Fig. 5b-h), nor the ATP levels in mitochondria (Fig. 5i), except the O/F ratio (Fig. 5f), suggesting a slight proton leakage. These parameters were independent of RTX activation during the time course of the experiments. To confirm the H^+^ leak induced by mitoTRPV1, mitochondrial respiration was assessed in permeabilized cells expressing mitoTRPV1 using the Oroboros-O2k device. This confirmed a limited increased respiration rate associated with a proton leak (O) while excluding any other change of mitochondrial respiration parameter (Extended Data Fig. 6).

**Figure 5.**
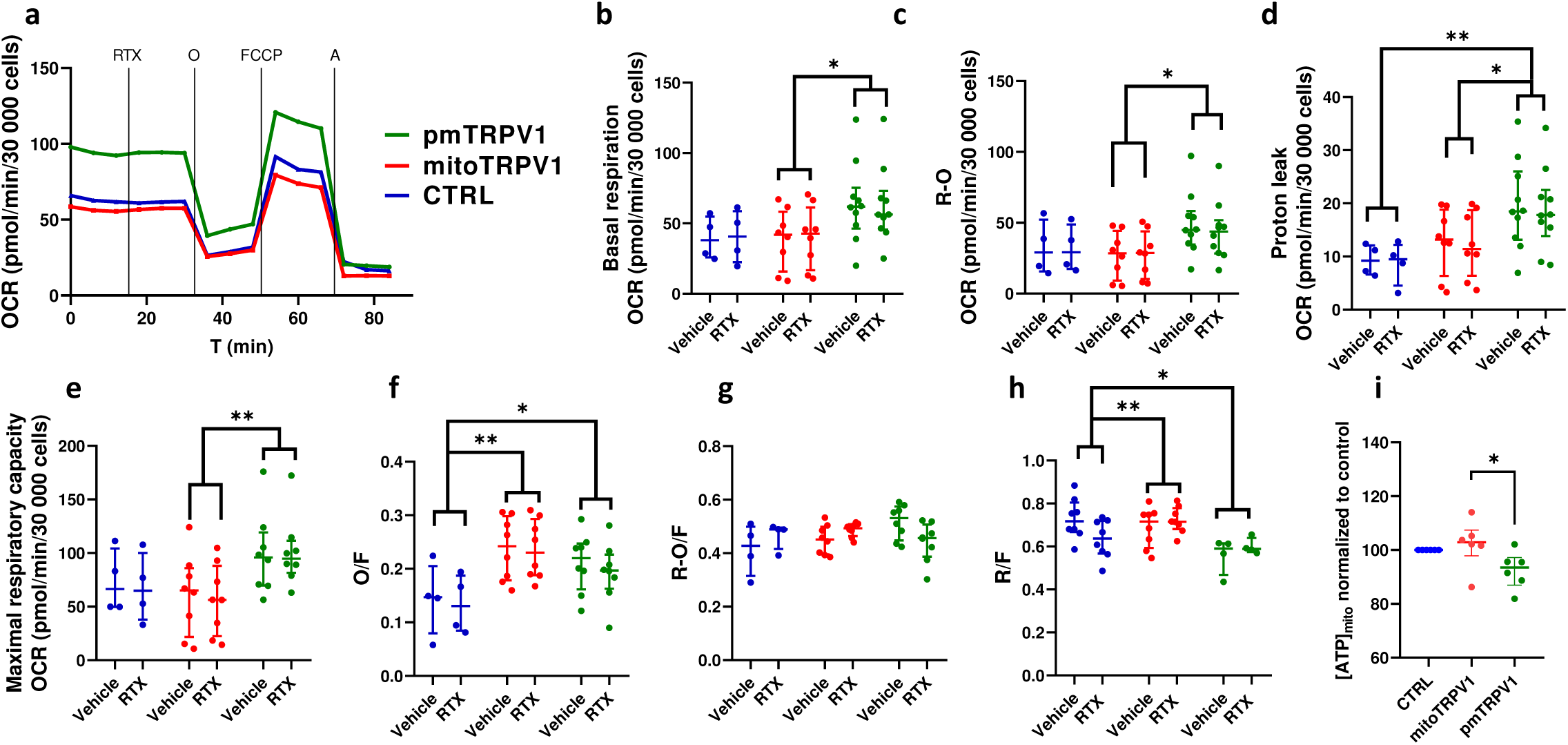
MitoTRPV1 activation does not affect mitochondrial respiration and ATP production. **a,** Typical profile of O_2_ consumption rates (OCR) in intact HEK293 cells expressing pmTRPV1 (green), mitoTRPV1 (red) or control cells (blue) in a Seahorse XF96 set-up. **b, c, d, e, f, g, h,:** Basal respiration rates (R, b), respiration linked to ATP production (R-O, c) respiration linked to proton leak (O, d), maximal respiratory capacity (F, e), the proportion of respiration linked to proton leak compared to maximal respiratory capacity (O/F, f), the proportion of respiration linked to ATP production compared to maximal respiratory capacity (R-O/F, g), the spare respiratory capacity (R/F, h) of HEK293 cells expressing pmTRPV1 (green), mitoTRPV1 (red) or control cells (blue), in the presence of the vehicle or RTX (1µM). **i,** Mitochondrial ATP levels were measured using the mitoATeam sensor and flow cytometry. Data are from at least three independent experiments. P-value correspondences *: p<0.05, **: p<0.01, ***: p<0.001, **** p<0.0001

Altogether, our experiments disclosed that the mitochondrial isoform of TRPV1 embedded in the inner membrane promotes Ca^2+^ efflux from mitochondria to cool down their temperature while maintaining the OXPHOS respiration and ATP levels.

## Discussion

Understanding mitochondrial thermoregulation is a crucial challenge for promoting the physiological maintenance of the mitochondrial network, its surrounding cells, and the whole organism. Recently, it was proposed that TRPV4 acts as a *“LOW”* mitochondrial thermostat, promoting Ca^2+^ entry into mitochondria to increase metabolism and thermogenesis^7^. Nevertheless, the identification of a *“HIGH”* thermostat that would limit mitochondrial thermogenesis remains an open question. Here, we propose that mitoTRPV1, a newly identified human TRPV1 isoform, could be this *HIGH* thermostat to prevent mitochondrial overheating, as we demonstrate that it complies with three key criteria: it localizes into mitochondria, it triggers a process preventing mitochondrial overheating after activation and it does not modify cell respiration and metabolism.

mitoTRPV1 is encoded by a yet uncharacterized TRPV1 mRNA variant consisting in a +1 overlapping ORF starting in exon1 and spanning exon2, which is reconnected to the consensual TRPV1 ORF by the maintenance of the small coding intron2. This molecular process leading to overlapping ORFs is unusual in the nuclear genome of higher eucaryotes while much more common in viral genomes and plastids^27,28^. The mitoTRPV1 mRNA variant, like most mRNA encoding mitochondrial proteins, is ubiquitously expressed in all human tissues, although with a 26 times lower abundance compared to the conventional TRPV1 mRNA. It encodes a 150 aa amino-terminal MTS cleaved after the import, which we demonstrated as mandatory and sufficient to drive TRPV1 core domains into the inner membrane, like most of the mitochondrial ion channels. Future studies aiming at identifying mitoTRPV1 orientation in the IMM will be vital in evaluating how the pH and ATP concentrations, two key mitochondrial parameters, could further regulate the channel activity and its temperature sensitivity. Importantly, we found that the original molecular process generating TRPV1 MTS is conserved only among mammals, while not in other vertebrate phyla, even birds that share homeothermy with mammals, supporting the concept that divergent molecular processes led separately to homeothermy in these two phyla^29,30^.

We then demonstrated that mitoTRPV1 acts as expected as a Ca^2+^ channel with basic pharmacological features similar to the conventional pmTRPV1 isoform: comparable RTX activation, EC_50,_ and capsazepine inhibition, although the two TRPV1 isoforms are embedded in membranes with different lipid compositions and protein concentrations. Assessment of Ca^2+^ homeostasis at the cellular level demonstrated that mitoTRPV1 activation drives a Ca^2+^ efflux from mitochondria to its close vicinity, paralleled by a simultaneous weak Ca^2+^ influx into mitochondria. These Ca^2+^ fluxes are compatible with the concept of futile Ca^2+^ cycle occurring between mitochondria and the endoplasmic reticulum^31^ possibly involving the SERCA, RYR/IP_3_R and MCU complexes in addition to mitoTRPV1^32^.

Ultimately, we confirmed that in the absence of mitoTRPV1, the mitochondrial temperature from HEK cells reach 51°C as previously observed^4^ whereas the expression of mitoTRPV1 restrains mitochondrial temperature increase to 44°C, a temperature compatible with the conventional TRPV1 isoform activation. Further preliminary extrinsic activation of mitoTRPV1 by RTX addition reduced the maximal mitochondrial temperature to 40°C, proving that mitoTRPV1 function and activation consist in preventing mitochondrial overheating. Surprisingly, the effects of mitoTRPV1 on the mitochondrial temperature had no simultaneous effect on mitochondrial respiration parameters, suggesting that mitoTRPV1 promotes heat dissipation rather than an inhibition of the Krebs cycle and OXPHOS activities. This is consistent with the concept that during chronic mitochondrial solicitation generating high mitochondrial temperature, mitoTRPV1 would promote a Ca^2+^ futile cycle, which extrudes “hot Ca^2+^” from mitochondria to cool down or exchange it in the endoplasmic/sarcoplasmic reticulum before re-importing fresh Ca^2+^ into mitochondria.

In this respect, the expression of the mitoTRPV1 isoform with the missense G684V mutation, which predisposes to exertional heat stroke^16^, annihilates both the mitochondrial Ca^2+^ efflux and the cooling process, establishing a factual correlation between Ca^2+^ transfer and mitochondrial heat dissipation. Thus, the futile Ca^2+^ cycle process triggered by mitoTRPV1 activation by high temperature would maintain high mitochondrial activity and ATP production while preventing mitochondrial deleterious overheating. This process is also supported by the physical properties of Ca^2+^ that displays a thermal conductivity 300 times higher than water [Ca^2+^: 200 W/mK (https://periodictable.com/Properties/A/ThermalConductivity.an.html, accessed on 18 september 2024) *vs* H_2_O: 0.6 W/mK, (https://medium.com/@thermtest/thermal-conductivity-of-water-61b19569b100, accessed on 18 September 2024), respectively]. Altogether, we propose that in mammals, the mitoTRPV1 isoform triggers a process responsible for mitochondrial heat dissipation, whenever the mitochondrial temperature reaches 44°C, by promoting a cooling Ca^2+^ circulation in between the mitochondria and the endoplasmic/sarcoplasmic reticulum. This is consistent with clinical and genetic data that evidenced pathological variants in the *RYR1*, *SERCA,* and *TRPV1* genes predisposing to life-threatening hyperthermia^15,16,33,34^, each variant would affect Ca^2+^ homeostasis and consequently mitochondrial cooling, ultimately leading to body overheating. Finally, the characterization of mitoTRPV1 isoform opens novel gates to identify pharmacological agonists and antagonists specific of each TRPV1 isoform, in order to better manage either nociception or body temperature in humans.

## Methods

### DNA analysis and MTS prediction

All TRPV1 protein and nucleotide sequences were collected from the NCBI database (https://www.ncbi.nlm.nih.gov/, visited the 04/08/2024) and analyzed with the MitoProt-II tool (https://ihg.helmholtz-munich.de/ihg/mitoprot.html^35^, visited the 04/08/2024) to predict the presence of a N-terminal mitochondrial targeting sequence (MTS) and its cleavage site. For the MTS prediction in animals for which no TRPV1 mRNA variant is described in protein databases, *TRPV1* gene sequences were collected from the NCBI DNA database and analyzed by a home-made algorithm screening all the open reading frames (ORF) for the presence of an MTS sequence in the first coding exons. The highest score was retained for each species and is shown in Fig.1f.

### Cloning

All primers are described in supplementary Table S1. Plasmid encoding pmTRPV1 (pcDNA5/FRT-TRPV1) was a gift from Aubin Penna (4CS labs - CNRS UMR 6041, Poitiers, France) and was used to clone mitoTRPV1 (pcDNA5/FRT-mitoTRPV1) using the Infusion Cloning method (Takara Bio). First, pcDNA5/FRT-TRPV1 was linearized by inverse PCR using A-for and B-for primers. Then, two synthetic complementary oligonucleotides corresponding to *TRPV1* intron2 were obtained from ThermoFischer and used to reconstruct the double-strand insert: B-for and B-rev. The insert was cloned using a vector/insert ratio 1/10 to generate an intermediate plasmid with an ORF coding for mitoTRPV1. Then, this plasmid was linearized by inverse PCR using the primers C-for and C-rev to delete the sequence located upstream of the mitoTRPV1 alternative start codon. The plasmid was re-circularized to obtain the final plasmid coding for mitoTRPV1 (pcDNA5/FRT-mitoTRPV1). The same method was used to construct the plasmid coding for mitoTRPV1-G684V mutant, using G684V-for and G684V-rev primers and the GFP, mCherry or 6X-His tagged versions of TRPV1, mitoTRPV1 or mitoTRPV1-G684V using D-for/D-rev, GFP-for/GFP-rev, mCherry-for/mCherry-rev and 6XHis-for/6XHis-rev primers. All plasmids were validated by Sanger sequencing.

### Cell culture and transfection

HEK293 and MCF7 cells were cultured in Dulbecco’s modified Eagle medium (Pan Biotech P0403500) with 10% fetal bovine serum (FBS), 4.5g/L glucose and 1mM L-Glutamine at 37°C and 5% CO_2_ and were routinely tested for Mycoplasma infections. The day before transfection, cells were seeded on different supports to obtain 70%-80% confluence 24 h later. Transfections were performed using Avalanche®-Omni (EZ Biosystems) or DharmaFECT (Horizon Discovery), and media were changed after 24 h. Cells were analyzed 48 h after the transfection. The transfection efficiency was measured using the TRPV1-GFP tagged transfection and Incucyte® Live-Cell Analysis (Sartorius) and routinely evaluated at 50% (Extended Data Fig. 1a).

Cell transfections were performed as previously described for individual cell Ca^2+^ imaging and ATP measurements^36^. Cells were transfected with a previously incubated transfection mix that contained 500 µL of serum-free DMEM, 2 µL of Dharmafect Duo, and 2 µg of DNA plasmid (either 2 µg for the mCherry-tagged plasmid, or a mix of 1 µg for Ca^2+^ or ATP genetic probe (erGAP1, CMV-mito-R-GECO1, 4mtD3cpv, N33D3cpv, or mitoAteam) and 1 µg for the mCherry-tagged plasmid). pcDNA-4mtD3cpv was a gift from Amy Palmer and Roger Tsien)^37^ (Addgene plasmid # 36324; http://n2t.net/addgene:36324 (accessed on 8 September 2023); RRID: Addgene_36324). erGAP1 plasmid was a gift from Maria Teresa Alonso^38^ (University of Valladolid, Valladolid, Spain). N33D3cpv was a gift from Yves Gouriou^39^. The vector expressing mitoATeam was kindly supplied by Hiroyuki Noji^40^.

### Reagents and Antibodies

Resiniferatoxin (RTX) was purchased from Alomone Labs. Capsaicin (CAP), capsazepine (CPZ), oligomycin, antimycin A and (4-trifluorométhoxyphénylhydrazono)-mésoxalonitrile (FCCP) were purchased from Sigma-Aldrich.

The following commercial antibodies were used: GFP-antibody (JL-8, Takara Bio) dilution 1/1000, 6X-His (MA121315, ThermoFischer) dilution 1/1000, α-tubuline (ab52866, Abcam) dilution 1/2000, TOM20 (ab78547, Abcam) dilution 1/2000, TIM23 (ab116329, Abcam) dilution 1/2000, SERCA (ab2817, Abcam) dilution 1/2000, TFAM (ab155240, Abcam) dilution 1/2000, VDAC (ab14734, Abcam) dilution 1/2000, GRP75 (ab2799, Abcam) dilution 1/2000. Additionally, two homemade TRPV1 antibodies targeting the C-terminal part of TRPV1 or MTS part of mitoTRPV1 were generated by Genosphere Biotechnology.

### Fluorescent microscopy live imaging

Cells were seeded and transfected in a 4-well µ-slide 4 (Ibidi). For live imaging, cells were loaded with 50 nM Mitotracker Red CMXRos (Thermo Fischer) for 15 min at 37°C. After rinsing with fresh medium, µ-slide were placed under a microscope NIKON ECLIPSE Ti-E (Nikon Instruments Europe) equipped with a camera Andor NEO sCOMS. Image acquisition and analysis were performed with Metamorph 7.7 software (Molecular device). 30 image planes were acquired along the Z-axis at 0.1 μm increments, and images were iteratively deconvolved using Huygens Essential® software (Scientific Volume Imaging, Hilversum, The Netherlands).

### Mitochondria isolation

All cell fractionation experiments were performed according to a method already described^41^. Briefly, mitochondria were isolated by suspending cells in an ice-cold isolation buffer (Mannitol 225mM, Sucrose 75mM, HEPES 10mM, EDTA 10mM, DTT1 mM). Cells were broken with a glass/Teflon Potter homogenizer by 100 movements. Cell suspensions were centrifuged at 800 g for 5 min at 4 °C. The supernatant was centrifuged at 6000 g for 10 min at 4 °C. The mitochondria pellets were re-suspended in an isolation buffer and used for proteinase K protection assay, Western blot, or subsequent centrifugations for Mitochondrial Associated Membranes (MAMs) isolation.

### Proteinase K protection assay

Mitochondria from HEK293 cells expressing mitoTRPV-6X-His were isolated and treated with increasing concentrations of digitonin (0%–0.15%) and a constant concentration of Proteinase-K (ThermoFischer) at 50 µg/ml for 15 min at room temperature. Samples without proteinase-K or with 1% Triton X-100 served as controls. Proteinase-K was inactivated by adding 100 µM PMSF, followed by incubation with ice-cold 10% trichloroacetic acid (TCA) on ice for 15 min. After centrifugation (10.000 g, 15 min, 4°C), TCA precipitates were dissolved in an SDS-page loading buffer, and samples were analyzed by Western blot.

### Intracellular Ca^2+^ measurements

Cytosolic Ca^2+^ imaging experiments were carried out on populations of cells with a FlexStation^®^ 3 Benchtop Multi-Mode Microplate Reader. HEK293 expressing pmTRPV1 or mitoTRPV1 were plated at a density of 50,000 cells by well (96 wells black/transparent bottom plate) in culture medium. Twenty-four hours after plating, cells were incubated for 60 min at RT with 4 µM Fura-2 AM and Pluronic^®^-F127 acid (0.02%) in freshly prepared buffer composed of Hank’s Balanced Salt Solution (HBSS) supplemented (in mM): 2.5 CaCl_2_, 1 MgCl_2_ and 10 HEPES-K (pH 7.4). After washing, cells were incubated in the buffer for 60 min for a complete de-esterification of the dye. Plates were illuminated at 340 and 380 nm excitation wavelengths, and the fluorescence emission spectra were recorded at 510 nm. After a 30 s baseline, TRPV1 activators, including RTX, were automatically injected, and the fluorescence emission spectra were monitored for 320 s at an acquisition frequency of 0.25 Hz. All experiments were performed in triplicate at least twice. Data was analyzed using the SoftMax Pro 5.4.1 software (Molecular Devices, Sunnyvale, CA, USA).

For individual cell Ca^2+^ imaging, HEK293 cells were plated on glass coverslips (24 mm diameter) in a 6-well plate. For cytosolic Ca^2+^ measurements, cells transfected with mCherry-tagged plasmid (expressing pmTRPV1-mCherry or mitoTRPV1-mCherry) were incubated for 30 min at RT with 2 µM Fura-2 AM in calcium-containing buffer (CCB). CCB consists (in mM) of 140 NaCl, 5 KCl, 1 MgCl2, 10 HEPES, 10 glucose, and 2 CaCl2, adjusted to pH 7.4. For all other Ca^2+^ measurements, HEK293 cells were co-transfected as described in the section “Cell culture and transfection.” erGAP1, 4mtD3cpv, or N33D3cpv were used to measure ER, mitochondria, and mitochondrial surface hot spot Ca^2+^ concentrations, respectively. Experiments were performed at room temperature (RT) in CCB. Glass coverslips were mounted on a magnetic chamber (Chamlide) and placed on a DMI6000 inverted wide-field microscope (Leica Microsystems, Wetzlar, Germany). Images were acquired with an Orca-Flash 4.0 Scientific CMOS camera (Hamamatsu, Photonics, Shizuoka, Japan) using a 40X oil-immersion objective and a Lambda DG-4+ filter (Sutter Instruments, Novato, CA, USA). mCherry fluorescence was excited at 572/35 nm, Fura-2 AM at 340 and 380 nm, and their respective fluorescent emissions were measured at wavelength 610 nm and 510 nm, respectively. ErGAP1 was excited at 403 and 470 nm, and their respective fluorescence emissions were measured at 520nm wavelength. 4mtD3CPV and n33D1CPV were excited at a wavelength of 430 nm, and their emissions were collected at 480 nm and 530 nm. Images (1024 × 1024 pixels) were taken with 5 s (Fura-2 AM and erGAP1) or 2 s time intervals (N33D3cpv and 4mtD3cpv). Fluorescence ratios of mCherry-positive cells were analyzed with MetaFluor 6.3 (Universal Imaging) after removing background fluorescence.

All Ca^2+^ kinetics display fluorescence intensities as follows: Fx = F-F0 for single cell experiments, or Fx = F/F0 for cellular populations, where F corresponds to the measured fluorescence and F0 to the baseline fluorescence. Variations of Ca^2+^ concentrations are shown in response to RTX treatment as either area under the curve (AUC) or fluorescence peak (ΔF).

### Mitochondrial respiration with Seahorse^®^ XF96

Mitochondrial respiration of transfected cells was performed using the Seahorse XF96 device (Agilent), as described by^42^. Briefly, HEK cells transfected for 48 h were plated in a 96-well Seahorse plate at 30,000 cells per well in a culture medium. After 5 h, the culture medium was replaced by a non-buffered assay medium (pH=7.4) containing 4.5 g/L glucose and 1 mM L-Glutamine, and cells were incubated at 37 °C (0% CO_2_) for 1 h. Cell oxygen consumption was measured during drug injections. Injection #1 was for RTX or DSMO, injection #2 for oligomycin (2 µg/ml), injection #3 for FCCP (50 to 1000 nM), and injection #4 for antimycin-A (2 µg/ml). Data analyses were performed using the Wave software (Agilent). Cell numbering was used for data normalization.

### Mitochondrial respiration with Oroboros^®^ O2k

Mitochondrial oxygen consumption measurements were performed at 37 °C and atmospheric pressure using a high-resolution oxygraph (O2k, Oroboros^®^ Instrument, Innsbruck, Austria). Respiration rates on permeabilized cells were measured in respiratory buffer RB (10 mM KH_2_PO_4_, 300 mM mannitol, 10 mM KCl, 5 mM MgCl_2_, 0.5 mM EGTA, and 1 mg/ml serum albumin bovine, pH 7.4) using substrates of CI, CI + CII and CII as followed. First, state 2 (non-phosphorylating) respiration was measured after adding 2.5 mM pyruvate and 5 mM malate. Second, the CI-linked maximal phosphorylating respiration was stimulated by saturating ADP concentration (1.5 mM). Succinate (10 mM) was then added to measure the combined CI and CII-linked respiration. Rotenone (5 μM) was used to inhibit CI activity and obtain the maximal CII-linked respiration. Third, oligomycin (F0F1-ATP synthase inhibitor, 4 μg/ml) and FCCP (carbonyl cyanide *p*-trifluoromethoxyphenylhydrazone, a mitochondrial uncoupler, 1 μM) were sequentially added to ensure that cells were fully permeabilized. Finally, antimycin-A (2 μg/ml) was added to monitor the non-mitochondrial respiration.

### Mitochondrial ATP measurement with flow cytometry

All experiments were performed on a Fortessa X-20 cells analyzer instrument (BD Biosciences) equipped with 4 lasers: violet laser 405 nm, blue laser 488 nm, Yellow-green laser 561 nm, and red laser 640 nm. Ratiometric analysis of the mitochondrial ATP-sensitive FRET probe, mitoATeam^40^, was measured by excitation at 405 nm and 488 nm and emission at 525/50 nm and 530/30 nm for CFP and YFP, respectively. mCherry fluorescence was measured by excitation at 561 nm and emission at 610/20 nm. The gating of the HEK cells was based on two combined parameters. First, events were gated for singlets based on FSC-height (FSC-H) and FSC-area (FSC-A) (Extended Data Fig. 7a), then the removal of cellular debris due to the cell preparation was done by the threshold of population, based on both side scatter (SSC) and forward scatter (FSC) (Extended Data Fig. 7b). Second, the settings of each photo-multiplying tube (PMT) for each fluorescent channel were done using non-transfected versus transfected cells (Extended Data Fig. 7c-d). The ratio was calculated using the ratio of F_YFP_ (median fluorescence intensity) /F_CFP_ (median fluorescence intensity). A total of 1000-10000 events in triplicates were recorded. Data were analyzed using FlowJo^TM^ software v10.8 (BD Biosciences, San Jose, CA, USA).

### Mitochondrial temperature

Mitochondrial temperature measurements were performed using the Mito-Thermo-Yellow dye (MTY) as previously described^4^. Briefly, 48 h-transfected or non-transfected HEK293 cells were labeled with 100 nM MTY in a culture medium. After 30 min, cells were centrifugated at 1500 g for 5 min, and the pellet was washed in PBS. Then, cells were maintained as a concentrated pellet for 10 min at 37 °C to establish an anaerobiosis condition. The fluorescence (excitation 542 nm, emission 562 nm), the temperature of the medium in the cuvette, and the respiration of the intact cell suspension were simultaneously measured in a magnetically stirred, 37°C-thermostated 1mL-quartz cell, using a Xenius XC spectrofluorometer (SAFAS, Monaco). Potassium cyanide (KCN) was added to cells to inhibit mitochondrial respiration and thermogenesis when the maximal mitochondrial temperature was reached, i.e., when the MTY fluorescence was stable at its lowest level. Then, the temperature of the cell suspension medium was lowered to calibrate the MTY fluorescence. The mitochondrial temperature was then calculated from the difference between the minimal and the maximal MTY fluorescence, obtained with mitochondrial heat production linked to respiration and its inhibition with KCN, respectively. Cell medium cooling was used to calibrate the MTY fluorescence (Fig. 4a).

### Data analysis

Data are presented as median ± interquartile range. All statistical analyses were performed using GraphPad Prism 8 software. The non-parametric Mann-Whitney test was used to compare two groups. For multiple comparison, the Kruskal-Wallis test followed by Dunn’s post-hoc test or a two-way ANOVA followed by a Bonferroni post-hoc test were used when appropriate. Differences were considered significant when the p-value is less than 0,05.

## Captions of supplemental figures

**Figure S1.**
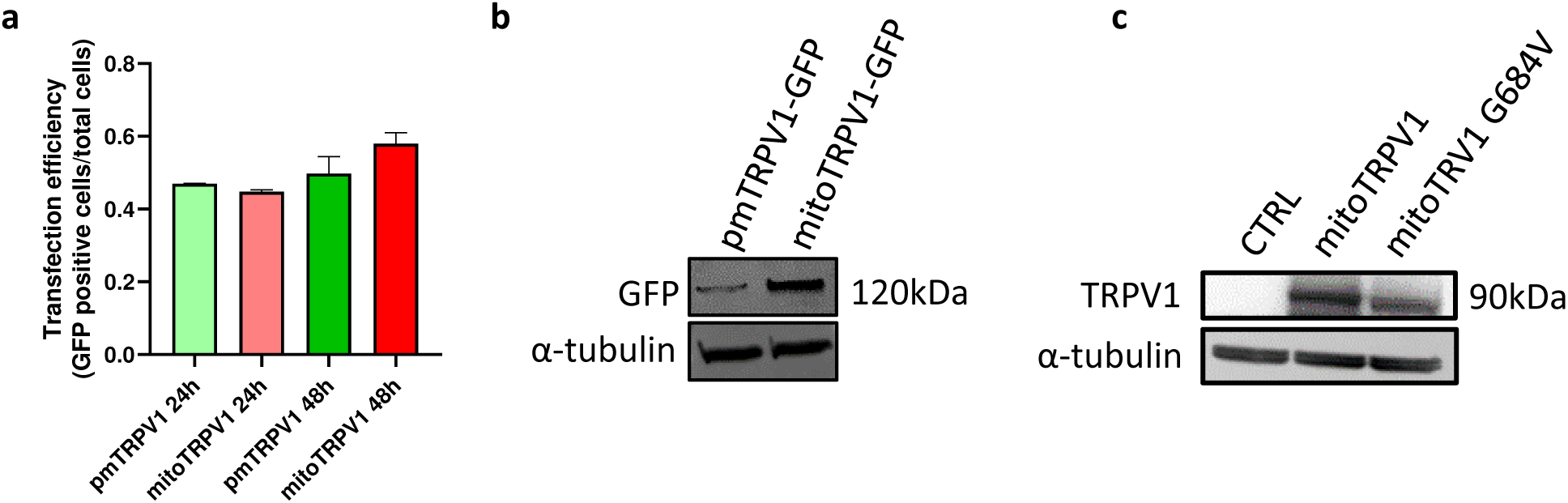
Assessment of the expression level of pmTRPV1, mitoTRPV1 and mitoTRPV1-G684V in HEK293 transfected cells. **a,** Transfection efficiency of mitoTRPV1-GFP and pmTRPV1-GFP at 24 h and 48 h (n=3). **b,** Western blot of cells expressing mitoTRPV1-GFP or pmTRPV1-GFP, using anti-GFP antibody and α-tubulin antibody, as a loading control. **c,** Western blot of cells expressing mitoTRPV1 or mitoTRPV1-G84V, using TRPV1 antibody and α-tubulin antibody, as a loading control.

**Figure S2.**
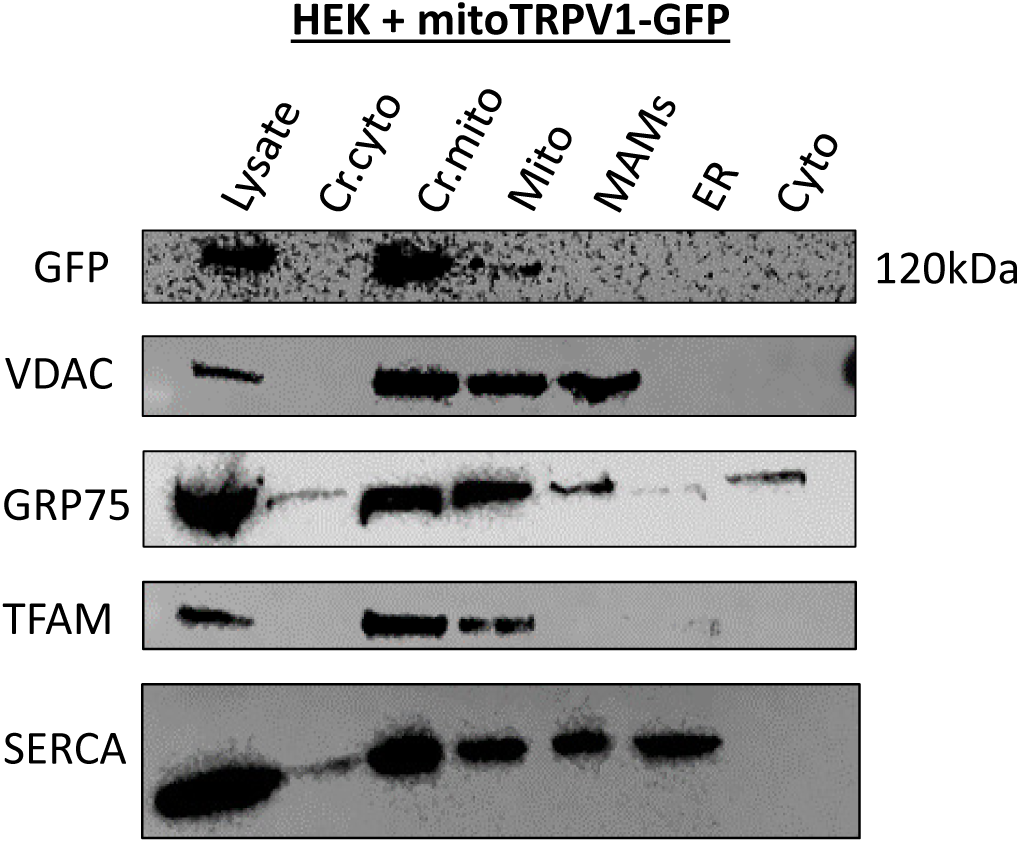
Cellular fractionation of HEK293 expressing mitoTRPV1-GFP. Cell fractions correspond to cell lysate (Lysate), the crude cytoplasmic fraction (Cr.cyto), which is composed of the cytoplasm (cyto) and endoplasmic reticulum (ER) fractions, the crude mito fraction (Cr.mito), which is composed of the mitochondria (Mito) and mitochondrial associated membrane (MAMs) fractions. MitoTRPV1-GFP is detected using an anti-GFP antibody. VDAC and GRP75 antibodies are used to detect MAMs, TFAM antibodies to detect mitochondria and SERCA antibodies to detect the ER.

**Figure S3.**
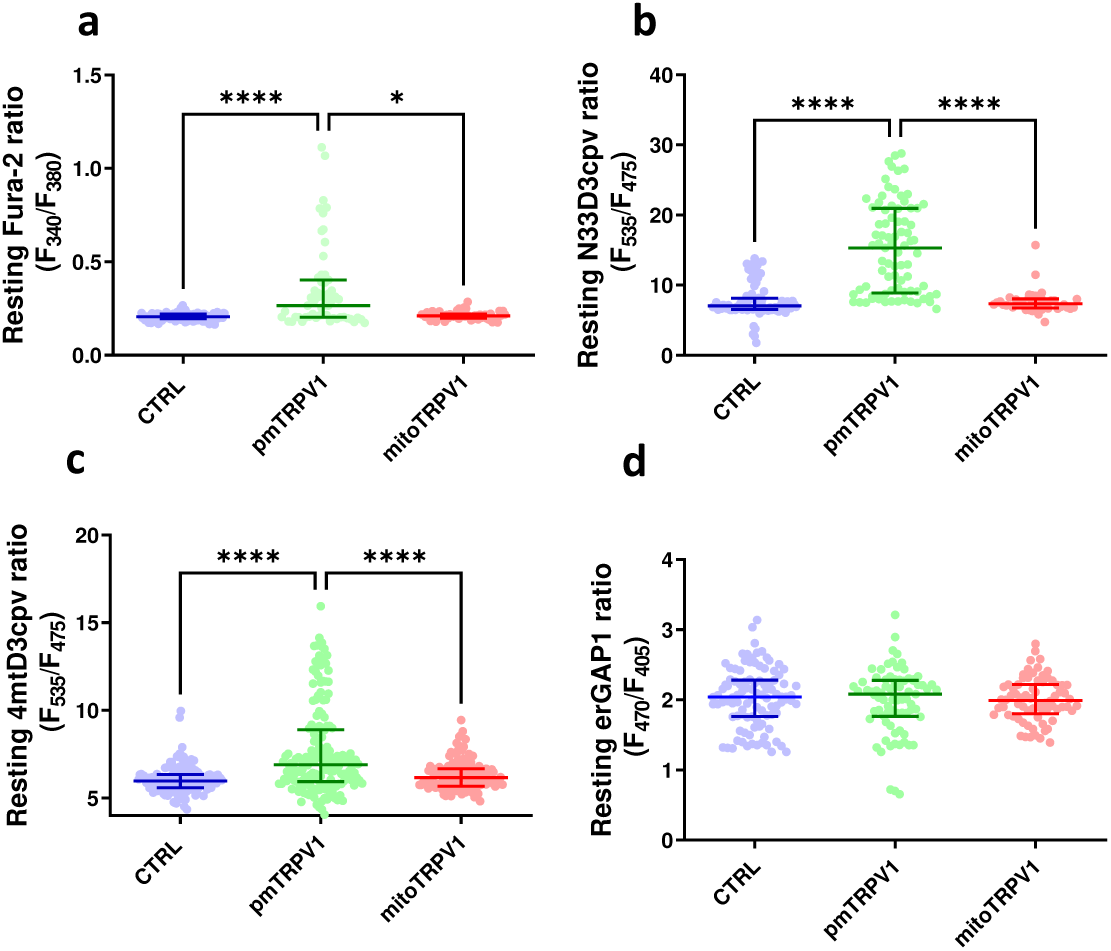
Resting Ca^2+^ concentrations in cellular compartments from HEK293 expressing mitoTRPV1 or pmTRPV1. **a, b, c, d:** Scatter plots of resting Ca^2+^ concentrations: in the cytoplasm measured with Fura-2 dye (a), in mitochondrial hot spot measured with N33D3cpv sensor (b), in mitochondria measured with 4mtD3cpv sensor (c), and in the ER measured with erGAP1 sensor (d) in CTRL cells (blue), mitoTRPV1 (red) and pmTRPV1 (green) expressing cells. Data are from at least three independent experiments. P-value correspondences *: p<0.05, **: p<0.01, ***: p<0.001, **** p<0.0001

**Figure S4.**
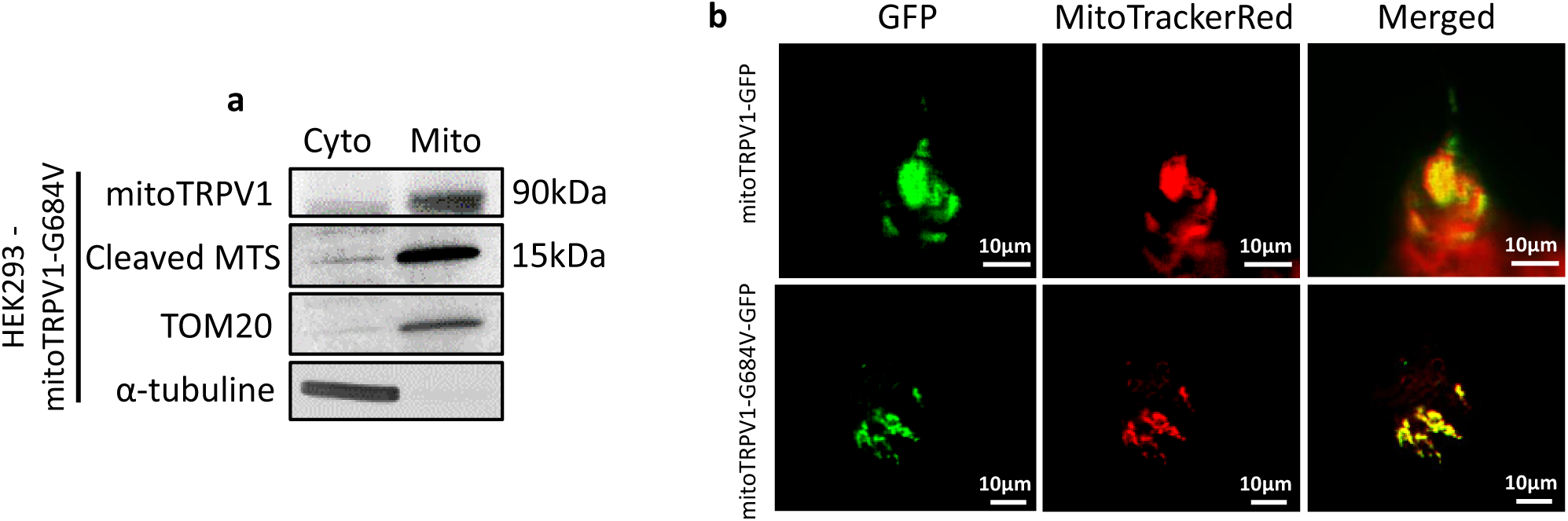
Expression level and mitochondrial localization of mitoTRPV1-G684V mutant isoform. **a,** Representative Western blot analysis of cytoplasmic (left) and mitochondrial (right) fractions of HEK293 cells expressing the mitoTRPV1-G684V variant, using mitoTRPV1 antibodies and mitoTRPV1 MTS antibodies. **b,** Co-localization analysis of GFP (green) and Mitotracker (red) signals in HEK293 cells expressing mitoTRPV1-GFP or mitoTRPV1-G684V-GFP.

**Figure S5.**
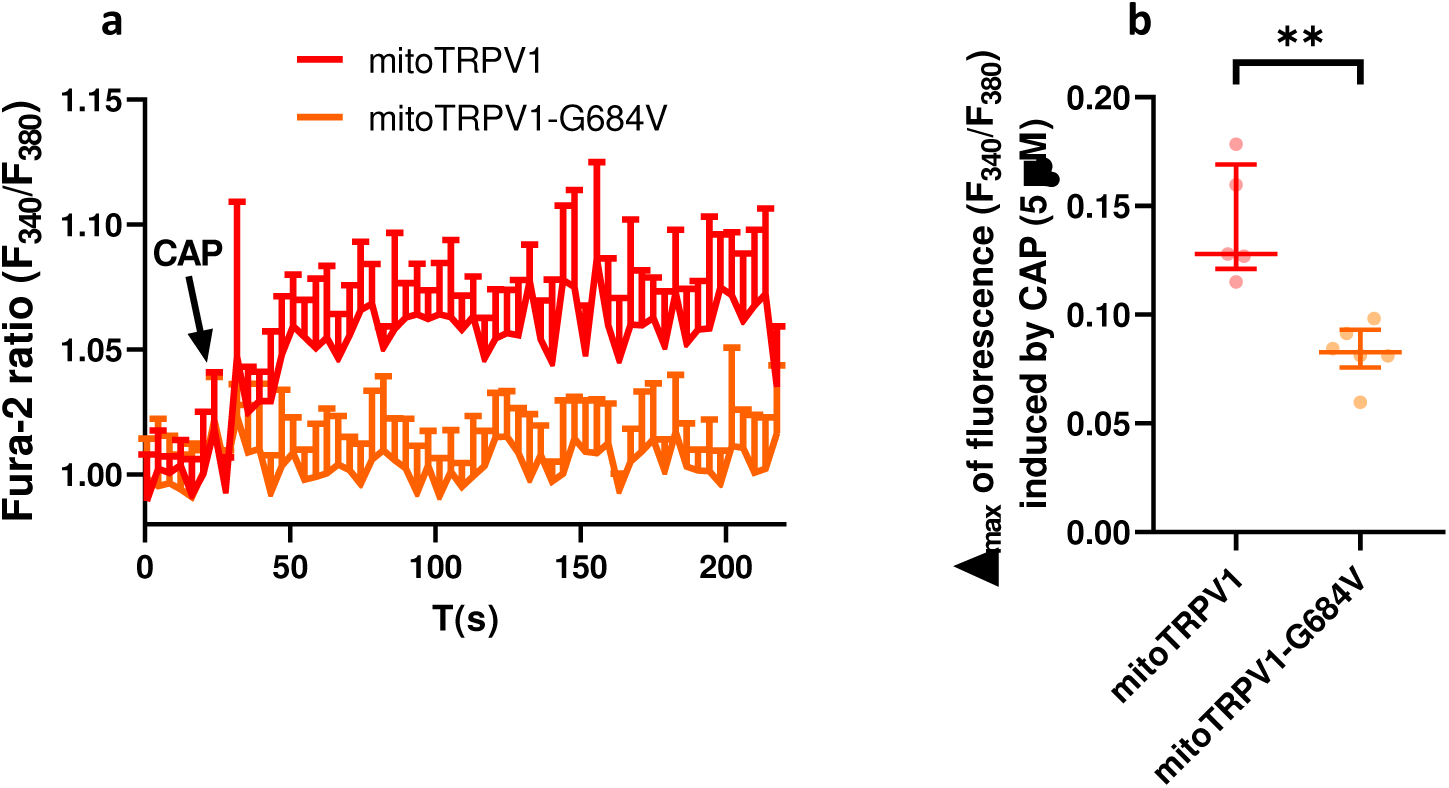
MitoTRPV1-G684V does not induce Ca^2+^ efflux from mitochondria. **a,** Cytoplasmic Ca^2+^ response of HEK293 expressing mitoTRPV1-G684 (orange) or mitoTRPV1 (red), after CAP (5µM) stimulation. **b,** Scatter plots representing the corresponding cytoplasmic Ca^2+^ maximal amplitude of CAP-response (n=3).

**Figure S6.**
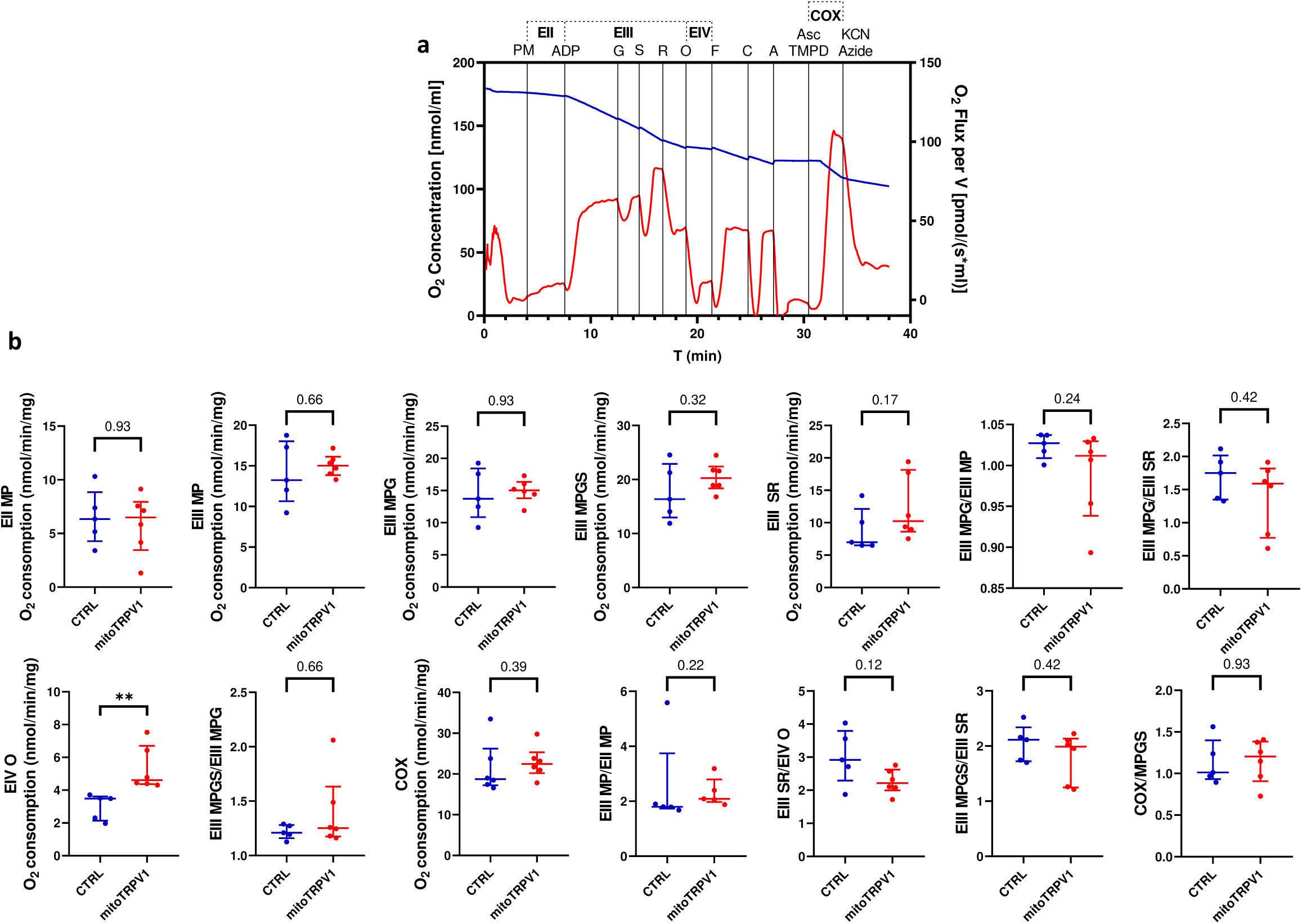
Impact of mitoTRPV1 expression on mitochondrial respiratory chain activity in permeabilized HEK293 cells. **a,** Representative trace of O_2_ consumption in permeabilized HEK293 cells. O_2_ concentration in blue and O_2_ flux in red. The addition of substrates and inhibitors are shown in black. Mitochondrial respiratory rates are measured after subsequent addition of 2.5 mM pyruvate and 5 mM malate (PM) (state II, complex I), 1.5 mM ADP and 5 mM glutamate (G) (state III, complex I), 10 mM succinate (S) (state III, complex I+II), 2.5 µM rotenone (R) (state III, complex II), 4 µg/ml oligomycin (O) (state IV, complex II). Cell permeabilization and mitochondrial membrane integrity are tested by the subsequent addition of 1 µM FCCP (F) and 8 µM cytochrome c (C). Cytochrome c oxidase maximal respiration is tested by the addition of 4mM ascorbate (Asc) and 0.3 mM N,N,N’N’-tétraméthyl-1,4-phénylènediamine (TMPD) followed by inhibition with 1 mM potassium cyanide (KCN) and 2 mM sodium azide (Azide). **b,** Scatter plots of parameters corresponding to the different respiration states of CTRL (blue) or mitoTRPV1 expressing (red) cells.

**Figure S7.**
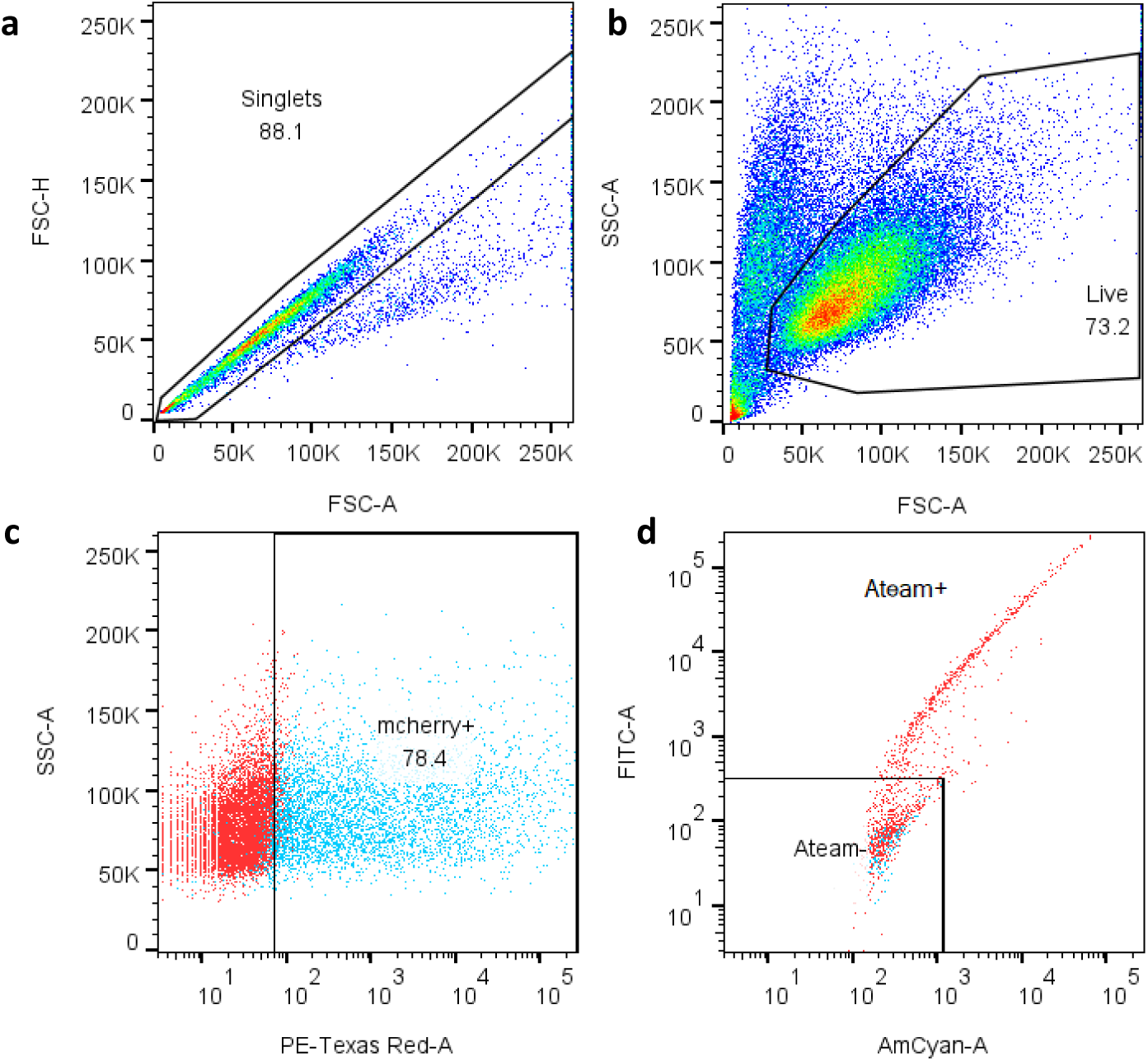
Gating strategies for measuring mitochondrial ATP levels by flow cytometry. Description of the standard strategies for the assessment of (**a**) singlets based on FSC-height (FSC-H) and FSC-area (FSC-A) and for (**b**) gating cells based on forward scatter (FSC) and side scatter (SSC). To analyze transfected cells, further gating is applied to the singlet population based on (**c**) mCherry (PE-Texas Red-A) and (**d**) CFP/YFP(AmCyan-A/FITC-A) fluorescence intensities.

**Table S1.**
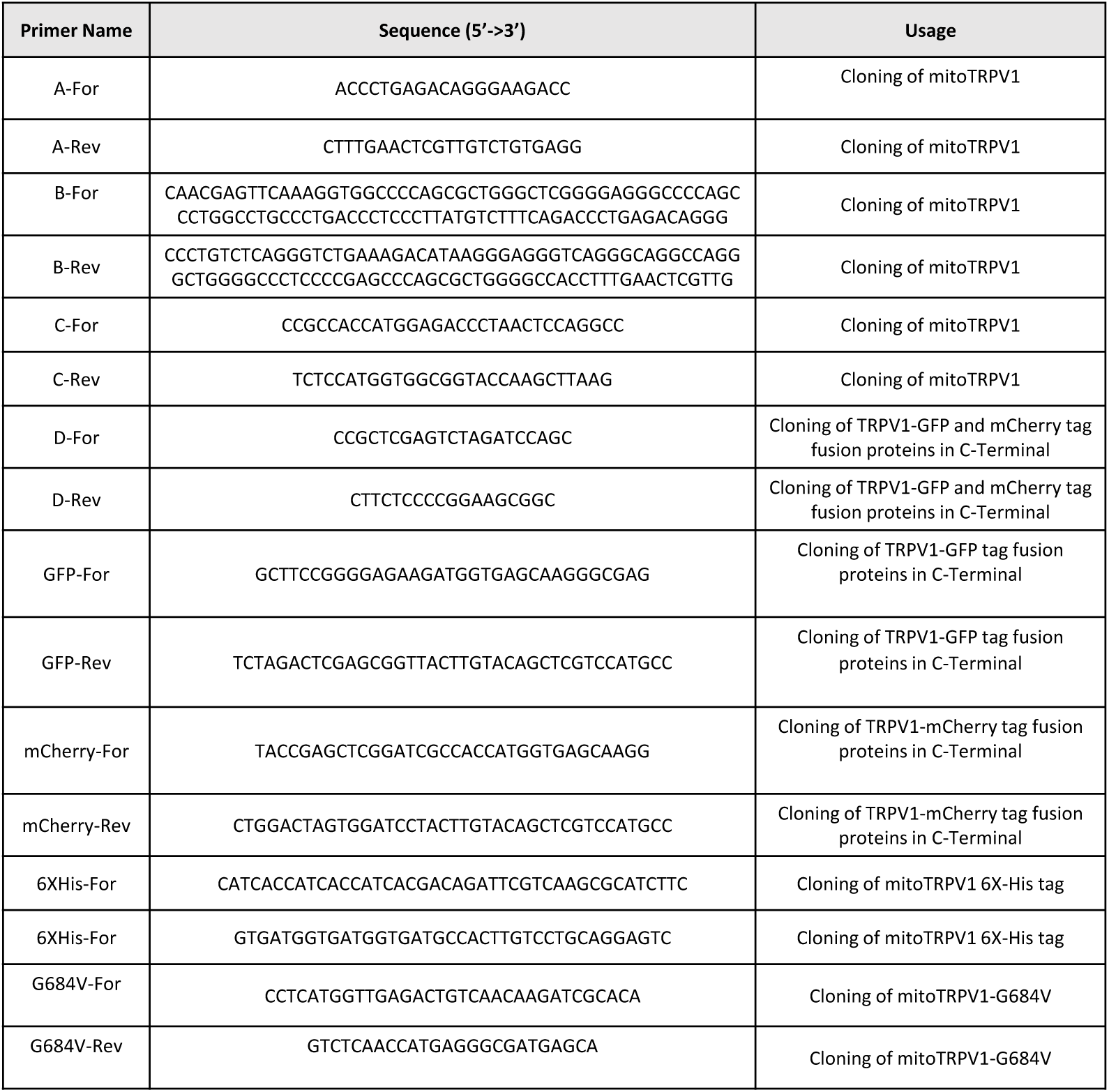
Table listing primers used to realize all the plasmids coding for mitoTRPV1 or pmTRPV1 tagged or mutants.

## Acknowledgements

We are indebted to Drs. Dominque Chretien, Paule Benit, Malgoratza Rak and Pierre Rustin for technical assistance and constructive discussions. We thank Aubin Penna (Laboratoire 4CS, Poitiers, France) for providing TRPV1 plasmid and Alexandra Malgoyre (Institut de Recherche Biomédicale des Armées, Bretigny-sur-Orge France) for her support. This work received financial support from the Inserm to the booster program Climate Change and Health and from the ASTRID program of the French National Research Agency N° ANR-21-ASTR-0010. This project received grants from the Région Pays de la Loire (France) and the French Direction Générale de l’Armement (DGA) with the support of Emmanuelle Guillot-Combe and Thierry Pauchard. We also thank University of Angers (France) for financial support.

## Notes

### Competing Interest Statement

The authors have declared no competing interest.

